# Whole Genome Sequencing of DENV-2 isolated from *Aedes aegypti* mosquitoes in Esmeraldas, Ecuador. Genomic epidemiology of genotype III Southern Asian-American in the country

**DOI:** 10.1101/2024.02.06.579255

**Authors:** Andrés Carrazco-Montalvo, Diana Gutiérrez-Pallo, Valentina Arévalo, Patricio Ponce, Cristina Rodríguez-Polit, Damaris Alarcón, Gabriela Echeverría-Garcés, Josefina Coloma, Victoria Nipaz, Varsovia Cevallos

## Abstract

Ecuador is a tropical country reporting Dengue virus (DENV) outbreaks with areas of hyperendemic viral transmission. Entomo-virological surveillance and monitoring effort conducted in the Northwestern border province of Esmeraldas in April 2022, five pools of female *Aedes aegypti* mosquitoes from a rural community tested positive for DENV serotype 2 by RT-qPCR. One pool was sequenced by Illumina MiSeq, and it corresponded to genotype III Southern Asian-American. Comparison with other genomes revealed genetic similarity to a human DENV genome sequenced in 2021, also from Esmeraldas. Potential introduction events to the country could have originated from Colombia, considering the vicinity of the collection sites to the neighboring country and high human movement. The inclusion of genomic information complements entomo-virological surveillance, providing valuable insights into genetic variants. This contribution enhances our understanding of Dengue virus (DENV) epidemiology in rural areas and guides evidence-based decisions for surveillance and interventions.

**Author Summary:** In this study, the complete genetic information of the dengue virus isolated from the mosquito vector *Aedes aegypti* was characterized using molecular biology, sequencing, and bioinformatics analysis. The isolation of the virus from the vector has contributed significantly to our understanding of the transmission dynamics of the disease and its genetic relationship with human cases. Identifying the serotypes and genotypes circulating in Ecuador has enabled us to comprehend the evolution of the virus from 2014 to 2023. Therefore, surveillance of genetic variability would aid in adapting precise prevention and control strategies to the genetic variants present in a specific geographical area. Entomo-virological surveillance is essential for assessing changes in risk and the impact of control measures.

## Introduction

Approximately half of the world’s population is at risk of infection with dengue virus (DENV). This mosquito-transmitted virus is one of the most significant and rapidly spreading pathogens worldwide. Over the past 50 years, the number of dengue cases has increased more than 30-fold, and according to some authors it is now considered an endemic disease in about 125 countries [1]. In the Americas, dengue appears in outbreak cycles dominated by one or two serotypes every 3 to 5 years. Notably, in 2019, a substantial peak was observed, with reported cases exceeding 3.1 million. Among these, 28,203 cases were severe, resulting in 1,773 deaths [2]. Ecuador has experienced a consistent rise of arbovirus related infections, particularly DENV [3]. Specific areas within the country have reached hyperendemic viral transmission, and new rural and remote areas have significant outbreaks [4]. Furthermore, severe forms of dengue, which were previously non-existent, are now becoming more prevalent [3]. Clinical signs of the disease can range from asymptomatic infections to mild or moderate febrile cases. However, there is also, a severe variant of dengue that poses life threatening danger characterized by plasma loss and hemorrhagic manifestations [5,6]. In 2022, there were 16,017 confirmed cases in the country, with 1,775 showing warning signs (11.1%), and 109 severe dengue cases (0.7%). As of epidemiological week 46, in 2023, there have been 24,637 cases of dengue mostly associated with dengue fever without warning signs [7].

Each of the four dengue serotypes comprises different genotypes classified according to the envelope’s phylogeny [8]. In the case of DENV1, it includes genotypes I, II, III (sylvatic), IV, V, and VI. For DENV2, it includes the genotypes: Asian-I, Asian-II, Asian/American, American, Cosmopolitan, and sylvatic genotype. DENV3, includes genotypes I, II, III, IV, and V. Finally, DENV4, has genotypes I, IIA, IIB, III, and sylvatic genotypes [1,9]. The four main serotypes of DENV share approximately 70% of nucleotide identity, but they are antigenically distinct [10], thus, infection with one serotype confers protection mostly to homologous viruses. The different outcomes of dengue fever mostly stem from the interaction of these four serotypes with previous immunity. Cross reactive, partially neutralizing immunity can be detrimental, increasing the risk of severe dengue in secondary heterologous DENV infections [11,12,13].

The transmission of dengue in rural areas is an emerging and complex process attributed to population expansion, connectivity, human movement that allows importation of infected individuals. Some evidence shows increasing seroprevalence in children from rural areas and increasing incidence over time [14].

In this context, the purpose of our study is to report the complete genome sequence of the DENV-2 virus detected in an *Aedes aegypti* mosquito pool, its genotype, and viral evolution in Ecuador.

## Methods

### Study Site

As part of A2CARES, an active surveillance network and cohort study of arboviruses, epidemiological surveillance and research are conducted in Ecuador by our collaborating team (UC Berkeley, CIREV, USFQ). In April 2022, the Center for Research on Vector-borne and Infectious Diseases (CIREV) of the National Institute for Public Health Research (INSPI) conducted entomological sampling in Borbon locality, Eloy Alfaro canton, Esmeraldas province, from Ecuador. Borbon is a commercial area with semi-urban characteristics (Fig 1).

**Fig. 1.**
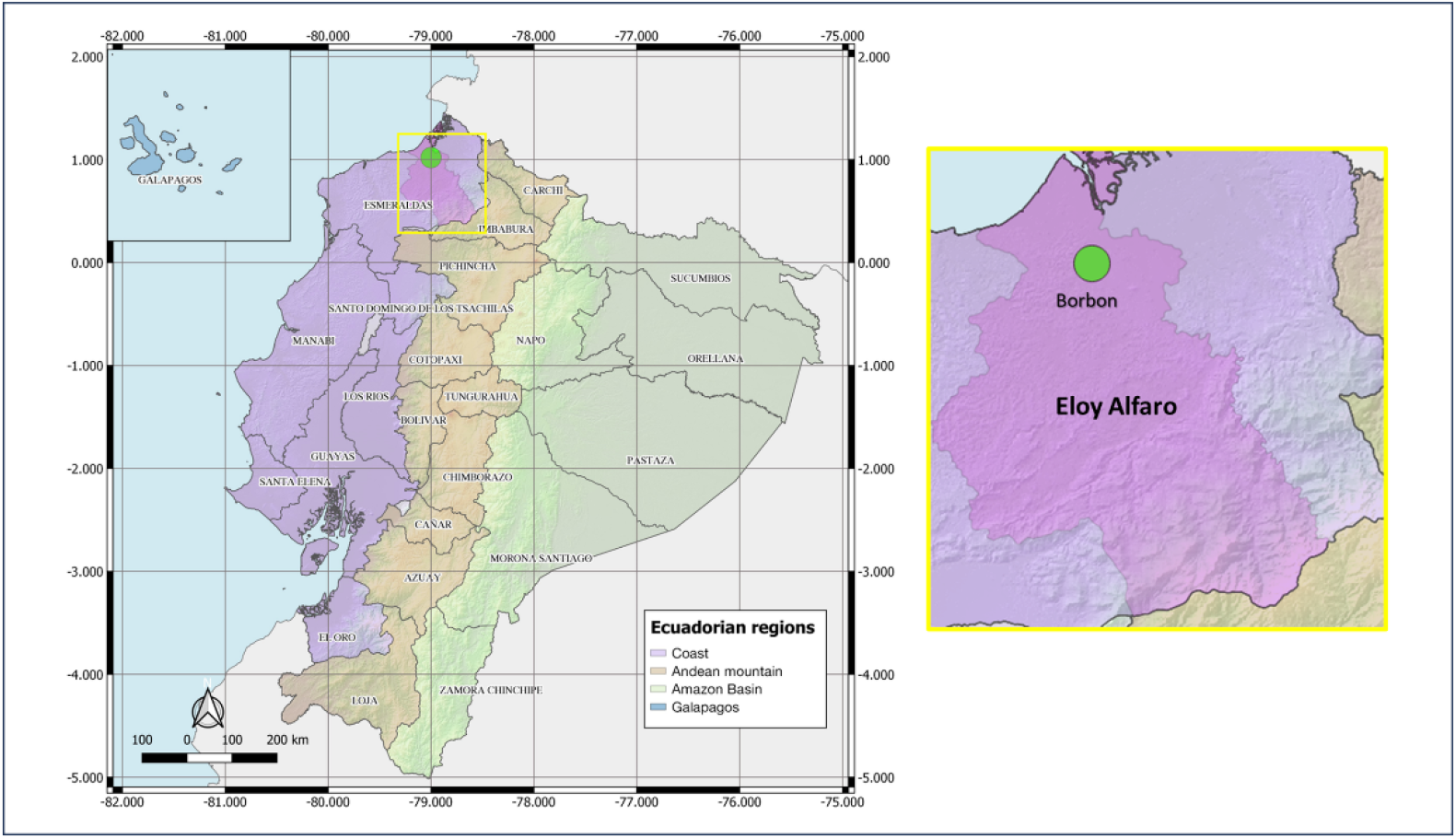
Map of entomological sampling location during 2021-2022. Country: Ecuador; Province: Esmeraldas; Canton: Eloy Alfaro, Borbon. QGIS 3.10.0.

### Sample collection

Adult *Aedes aegypti* mosquitoes were collected using a ProkoPack™ aspirator for intra-domiciliary aspirations during the morning, noon, and afternoon in 300 homes in Borbon. Homes were selected at randomly. Adult *Aedes* mosquitoes were identified on-site following Rueda’s (2004) pictorial key. Females identified as *Aedes aegypti* (fed or unfed) were placed in RNAlater® in pools of 10 to 30 individuals considering the collection hour and the closeness of the homes. The pools were then transported to the Molecular Biology Laboratory of the CIREV-INSPI located in Quito, Ecuador, for processing.

### RNA Extraction and RT-qPCR

A total of 155 *Ae. aegypti* females organized in 13 pools were used for RNA extractions. Each pool was initially macerated with micro-pistils (Fisher Scientific, USA) in liquid nitrogen. RNA was extracted using the RNeasy Mini Kit (Qiagen, Hilden, Germany), following the manufacturer’s instructions.

The ZDC Multiplex RT-PCR Assay Kit (Bio-Rad, USA) was used to detect the presence of Zika, Dengue, and Chikungunya viral RNA simultaneously; positive controls were included for each virus. The DENV-positive pools were tested using the CDC Real Time RT-PCR assay for detection and typing of DENV; positive control included four DENV serotypes isolated in cell culture [15].

### Sequencing and genome assembly

Sequencing and bioinformatics analyses were conducted at the National Reference Center for Genomics, Sequencing, and Bioinformatics CRN-GENSBIO at INSPI. The COVIDSeq Assay amplicon (Illumina®, USA) sequencing protocol was followed. Specific primers designed for DENV-2 were used (S1 Table). These primers enabled the amplification of genome fragments in the range of 400 to 600 base pairs (bp) through PCR amplification. The amplified fragments were subsequently sequenced using the Illumina MiSeq platform.

Three bioinformatics workflows were used for raw processing. FASTQ reads, including: quality analysis, read trimming, reference mapping (NC_001474.2), and genome assembly. Two of the workflows used similar tools, such as fastQC, Trimmomatic, BWA or Bowtie2, and Ivar consensus. The third workflow followed the ViralFlow protocol [16]. The sequenced sample was uploaded to the GISAID EpiArbo database with accession number EPI_ISL_18195022.

### Genotyping and phylogenetics

Genotyping analysis was performed using the Dengue Virus Typing Tool. This method uses the envelope glycoprotein (E) gene (1.485 bp) and whole genomes corresponding to serotypes: DENV-1 (I, II, III, IV and V genotypes), DENV-2 (I American, II Cosmopolitan, III Southern Asian-American, IV Asian II, V Asian I and VI Sylvatic genotypes), DENV-3 (I, II, III and 3 genotypes), and DENV-4 (4I, 4II, 4III and 4IV genotypes). After assigning the serotype, phylogenetic analysis by Maximum Likelihood, using the same tool, was completed to determine the genotype [17]. In addition, Multiple Sequence Alignment Viewer (MSA) analysis was performed using BLAST in GenBank to search for similarities with other genomes in this database.

All genomes belonging to the DENV-2 genotype III Southern Asian-American reported for Ecuador from 2014 to 2023, were selected and downloaded from GISAID EpiArbo database (https://gisaid.org/), and the reference genome NC_001474.2 was used as an outgroup. The dataset was aligned using MAFFT v7.453 [18] and then refined using AliView v1.28 [19]. Using Augur, sequences were indexed (with metadata containing dates and locations), filtered, aligned, and inferred through a Maximum Likelihood tree with 1,000 bootstrap replicates, molecular clocks and refinement, and the results were exported for display in auspice [20]. Each resulting tree was formatted in Figtree v1.4.4 [21]. The sequence of DENV-2 isolated from *Ae. aegypti* was analyzed, and a comparative analysis was conducted with sequences from human cases published in GISAID [22].

## Results

### Entomology and DENV serotyping

Mosquito samples were collected on April, 2022 that corresponds to the wet season in the collection locality. Five pools (5/13) were positive for DENV-2, with obtained CT values between 16.13 and 37.01. We decided to sequenced the pool positive for DENV-2 with CT value = 16.13; the other four positive pools had CT greater than 25. The selected pool comprised 30 *Aedes aegypti* mosquitoes (13 unfed and 17 fed) collected in the Borbon locality (-78,989769; 1,087634) on April 27, 2022 (S2 Table).

### Genome assembly and genotyping

Using different workflows, the entire DENV-2 genome was assembled with 1891 Ns of 10723 base pairs. A comparison between the obtained genome and reference genome NC_001474.2 is shown in Fig 2a. The mosquito sample analyzed (EPI_ISL_18195022) was assigned to Genotype III Southern Asian-American (Fig 2b). The MSA Viewer in the BLAST analysis revealed a close relationship with genomes posted in GenBank from Venezuela and Colombia (S1 Fig), similar to those reported from human samples from the same area.

**Fig. 2.**
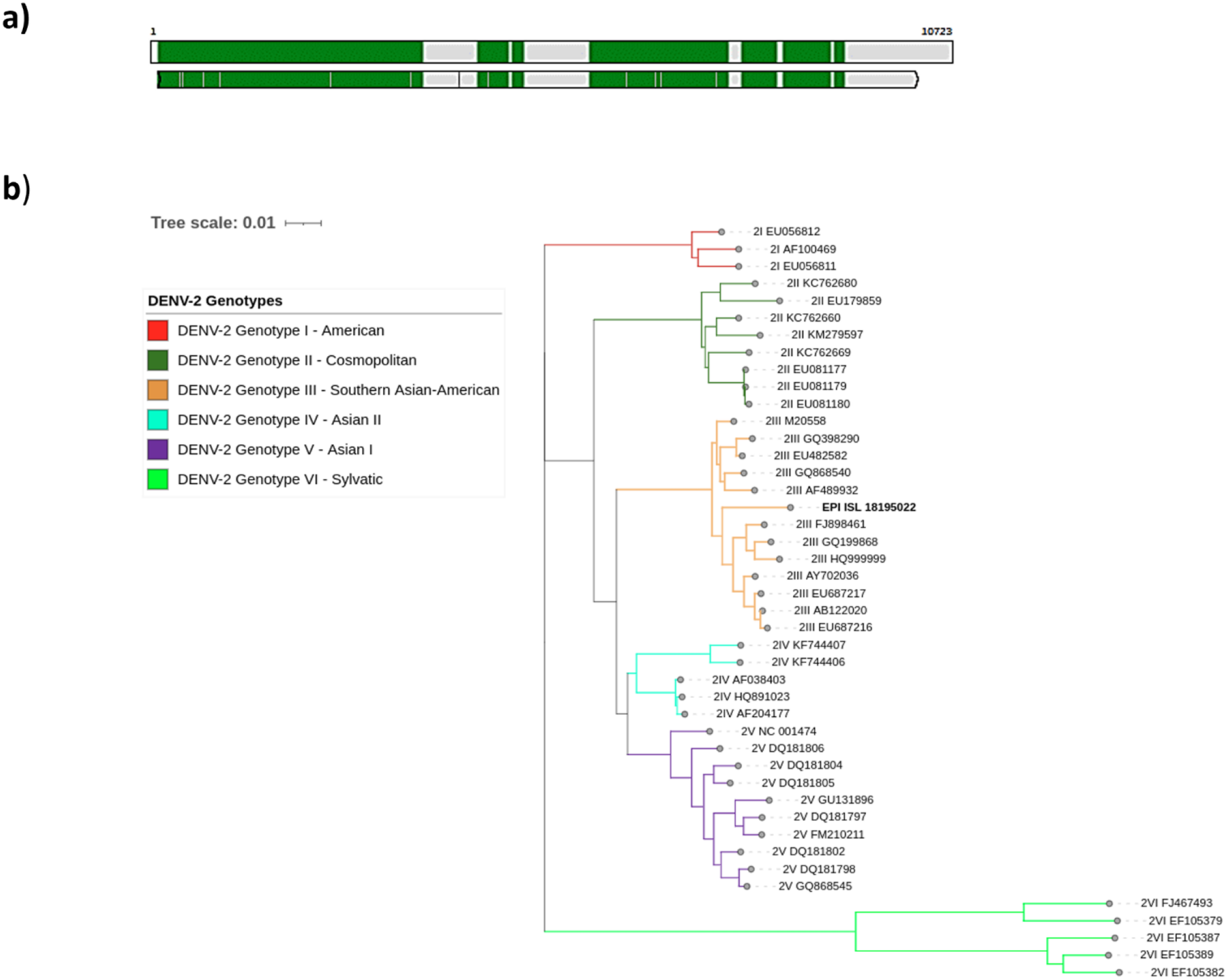
Genome assembly sequenced in this study and genotype assignment. **a)** Sample coverage of the sequenced sample EPI_ISL_18195022 against the reference genome NC_001474.2. **b)** Phylogenetic tree of the DENV-2 genotypes corresponding to the analyzed sample mark with bold (EPI_ISL_18195022). Each genotype is represented by a different color: I - American (red), II - Cosmopolitan (dark green), III - Southern Asian-American (orange), IV – Asian II (light blue), V - Asian I (purple), and VI - Sylvatic (light green).

### Phylogenetics Analysis of DENV from Esmeraldas

We downloaded 73 genomes of DENV-2 genotype III Southern Asian-American from GISAID, including our sequence (EPI_ISL_18195022), and selected sequences from human samples from the same study area, along with the sequence NC_001474.2. In the phylogenetic tree (Fig 3), the sequences collected during the years 2014-2015, labeled in black (without locality assignment), formed an ancestral clade with respect to the rest of the samples. Another clade was formed from the sequences collected during 2020-2023. This recent clade includes samples collected in 10 provinces of Ecuador, with 14 samples belonging to the province of Esmeraldas in greenish yellow, from our complement project following human cases. The sample reported in this study (enclosed in a circle) appears to be the closest reported from Timbire locality with 87 bootstrap values, and establishes a genetic group with other samples from the nearby localities of Borbon, Santa Maria, and Colon Eloy, all in the rural Eloy Alfaro canton.

**Fig. 3.**
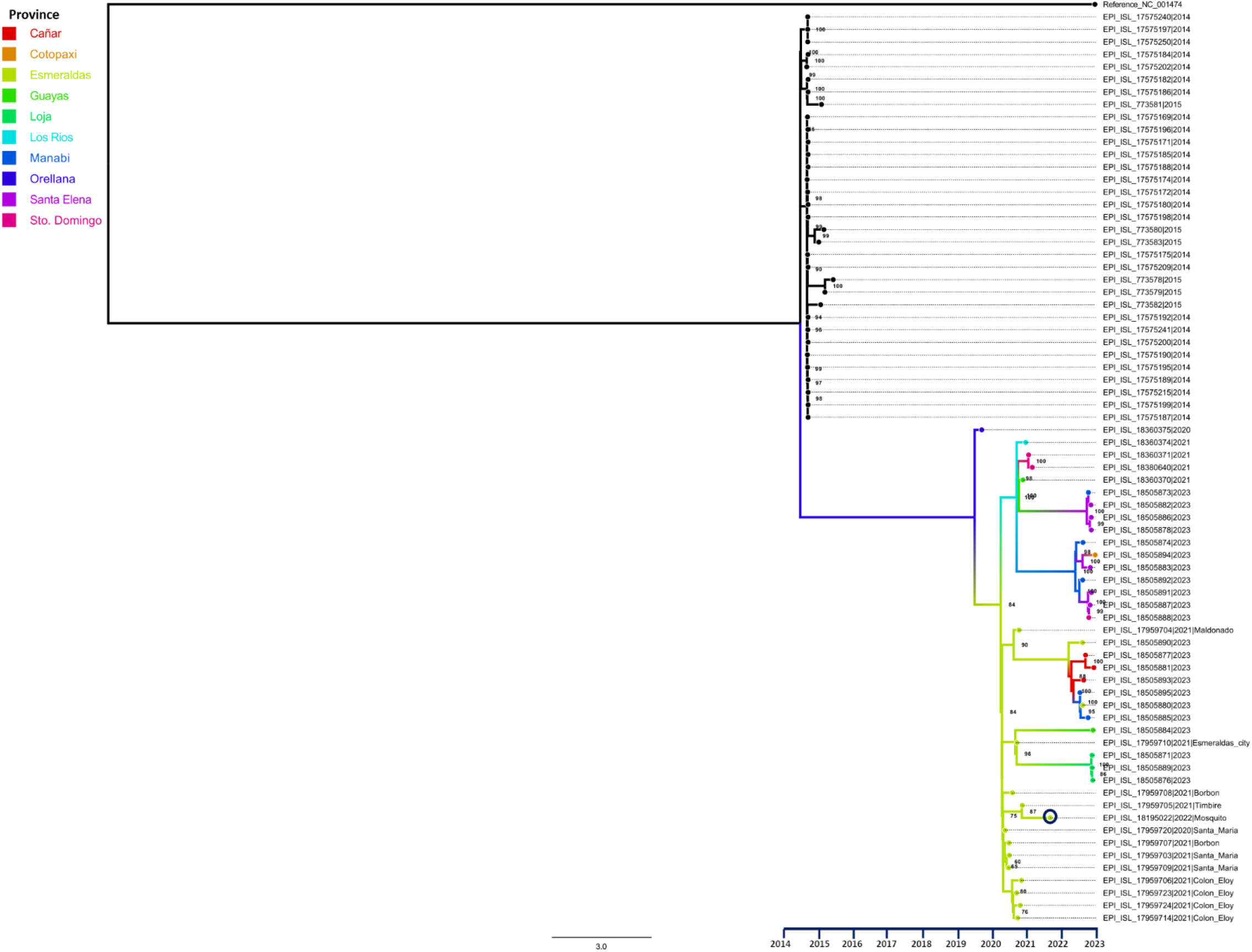
Phylogenetic tree of DENV-2 samples circulating in Ecuador. (Maximum likelihood Tree, Bootstrap: 1000). The sample reported in this study is enclosed in a circle.

## Discussion

Our entomo-virological surveillance study demonstrates that it is possible to isolate a DENV whole genome directly from pools of up to 30 mosquitoes.

This finding is consistent with the study conducted by de Figueiredo et al. 2010, [23]. Furthermore, the quality of the RNA allows molecular detection and the use of further deep sequencing protocols. The phylogenetic analysis shows that the DENV-2 isolated from *Aedes aegypti* belongs to genotype III Southern Asian-American. This genotype formed a cluster with genomes from Belize (FJ898461.1), Nicaragua (GQ199868.1), Guatemala (HQ999999.1), Cuba (AY702036.1), Puerto Rico (EU687217.1, EU687216.1), and the Dominican Republic (AB122020.1), as determined by the Dengue Virus Typing Tool for reference genome assignment. Additionally, BLAST identified a close relationship with genomes from Venezuela and Colombia. The phylogeographic study published by Allicock et al., in 2012 [24], analyzed 191 sequences of this genotype from DENV samples collected in various countries in the Americas between 1981 and 2008. It is estimated that this genotype entered the continent through the Greater Antilles in 1979, subsequently spreading to the Lesser Antilles between 1992 and 1995, and then to the South American coast [25].

The study from febrile case-patients during 2019-2021 published by Márquez et al., reports two possible introduction events of DENV-2 to Ecuador [22]. The first event involved a strain that arrived before 2009 and circulated until 2015. This strain was closely related to Colombian strains that circulated up to 2013. The second event involved a strain that arrived in 2013, resulting in an Ecuadorian lineage that still persists and shares a high degree of similarity with sequences from Colombia and Venezuela [22]. These studies with human samples are consistent with our results, as the isolated genome from the vector shows a greater similarity to genomes from Colombia and Venezuela, as reported from virus sequences from human serum from dengue cases from the same locality [22]. The reported DENV-2 genome from mosquitoes also forms a cluster with sequences from other countries in the Americas. The locality of Borbon in Northwestern Ecuador is located 117 kilometers from Tumaco (Nariño, Colombia) by land, and it also has connections through rivers and sea following the Borbon-Limones-Tumaco (Colombia) route. Therefore, the circulation of the same genotype and a similar sequence in the area could be due to the commercial exchange and human migration that exists between both countries via the mentioned routes. The detection of DENV in mosquitoes show that the virus is actively circulating in rural communities after its importation to the area.

In an study by Waman, W., et al. 2016, an *in silico* research showed that the population that displays genotype III Southern Asian-American is divided into four main groups, corresponding to Asia, Central America, South America, and North America [8]. The group is divided into seven subpopulations or lineages that reveal further subdivision within the South American and Central American groups. Three distinct lineages (AA3-AA5) were identified within the group of South American strains. Lineage AA3 includes older strains from the United States, Brazil, Puerto Rico, and Venezuela. A different lineage, AA4, is formed by American strains that were isolated during the 1996-2008 period in the United States, Brazil, Venezuela, and Colombia. On the other hand, the AA5 lineage consists of modern strains from the USA, Brazil, the Dominican Republic, and Jamaica [8]. This finding is similar to the reported by Fritsch et al. (2023), who also reported DENV-2, genotype III – Southern Asian-American [26]. Furthermore, the geographical dispersion of DENV-2 serotype may be associated with various mechanisms used to evade or escape the immune response [27]. Phylogenetic analysis revealed that the sequences from Ecuador collected in 2020, 2021, 2022, and 2023 were organized in another clade, different from samples collected in 2014 and 2015, identifying evolutionary events and genetic changes in genotype III Southern Asian-American. Human population immunity, competence and vector capacity of mosquitoes, seasonal variations, and random stochastic events contribute to the evolution and genetic changes of DENV[28]. In addition, this virus acquires one mutation per replication cycle in a vertebrate, errors attributed to RNA-dependent RNA polymerase (RdRp), resulting in intrahost genetic variants. DENV evolution also occurs under strong negative selection pressure [29,30]

Borbon is the commercial hub in the area, connected by both land and river routes to other localities, including Colombia, which could lead to a greater genetic diversity of the virus. This genetic diversity may extend from Borbon to nearby town Santo Domingo and Santa Maria (localities interconnected by river routes) and from Borbon to Maldonado (17.4 km), Timbire (30.7 km), and Esmeraldas city (107 km) (localities interconnected by land routes). The sequences from human serum from dengue cases found in Borbon are similar to those within this clade and have been actively circulating in the area [30,31]. Due to human migration and high mutation rate, the dengue virus undergoes constant evolutionary changes, increasing the likelihood of new variants emerging, particularly in the study area where there is high mobility between the populations from Colombia, Ecuador and lately from Venezuela [32].

The implementation of Next-generation sequencing (NGS) technology in epidemiological contexts has enriched our understanding of DENV’s population dynamics, which are influenced by viral replication sites and serotype-specific immune responses. This, in turn, can impact disease outcomes and containment during an outbreak. However, the lack of knowledge about the relationship between disease and its genetic diversity is hindered by variations in the methods and definitions used to estimate diversity within the host [33–35]. Through the application of genomic surveillance, using Next-generation sequencing, we successfully determined the genotype, and complete genome of the dengue virus within a mosquito host. Being able to identify the circulating serotypes and genotypes is essential for understanding the virus’s genetic variability, anticipating potential severe outbreaks, and tailoring specific prevention and control strategies specific to the serotypes present in a particular geographic area. Entomo-virological surveillance is critical for assessing changes in risk and the impact of control measures. The importance of this study lies in the application of DENV sequencing to identify the differences between naturally occurring infection in a human population and in the vector *Aedes aegypti*.

## Acknowledgments

We would like to express our gratitude to Universidad San Francisco de Quito for providing access to the additional sequences utilized in this study, supporting the field logistics and to provide primers for sequencing DENV2, which greatly contributed to our research.

This study was funded by the National Institute of Allergy and Infectious Diseases, National Institute of Health, R01 AI132372-02, entitled: “Zika and Dengue Co-circulation Under Environmental Change and Urbanization,” 1U01AI151788 entitled “American and Asian Centers for Arboviral Research and Enhanced Surveillance-A2CARES-CREID” and Sustainable Sciences Institute.

## Supporting information

**S1 Table. Specific primers designed for sequencing DENV-2**. List of primers and sequences used for Illumina MiSeq sequencing.

**S2 Table. Sequenced positive pool data**. Information of female *Aedes aegypti* mosquitoes corresponding to the sequenced pool.

**S1 Fig. Phylogenetic tree generated in MSA to compare our sample with other countries**. The Ecuadorian sample reported in this study is genetically closer to samples collected from Colombia and Venezuela.

## Notes

### Competing Interest Statement

The authors have declared no competing interest.

